# Time-scale separation and stochasticity conspire to impact phenotypic dynamics in the canonical and inverted *Bacillus subtilis* core genetic regulation circuits

**DOI:** 10.1101/245795

**Authors:** Lijie Hao, Zhuoqin Yang, Marc Turcotte

## Abstract

In this work, we study two seemingly unrelated aspects of core genetic nonlinear dynamical control of the competence phenotype in Bacillus subtilis, a common Gram-positive bacterium living in the soil. We focus on hitherto unchartered aspects of the dynamics by exploring the effect of time scale separation between transcription and translation and, as well, the effect of intrinsic molecular stochasticity. We consider these aspects of regulatory control as two possible evolutionary handles. Hence, using theory and computations, we study how the onset of oscillations breaks the excitability-based competence phenotype in two topologically close evolutionary competing circuits: the canonical “wild-type” regulation circuit selected by Evolution and, the indirect-feedback inverted circuit that failed to be selected by Evolution, as was shown elsewhere, due to dynamical reasons.

## Introduction

Living systems are at times full of randomness thus unpredictable and, at other times, orderly and predictable. These competing viewpoints are far from being incompatible with each other and seem entrenched in the subtlety of not only how, but why life organizes in the way we observe or infer, the latter increasingly relying on theory. Which of these aspects is most important depends entirely on context that relates to the size of fluctuations. At the core of genetic control, there exists a fine interplay of intrinsic randomness with deterministic dynamics that deeply impacts phenotype. It is fair to say that, at the core genetic control level, living systems are neither fully predictable nor fully random, but in fact reflect both aspects. The functional role of biochemical fluctuations - noise- in genetic circuits is reviewed in (1). Here, to explore these aspects further, but in an expanded scope that includes dynamical time-scales, we focus on the canonical Evolution selected *Bacillus subtilis* bacterium control circuit and on an inverted mutant circuit that Evolution did not favor (2). The core regulatory gene circuits that underlie the behavior of these organisms are shown on Figure 1. While these systems have been theoretically (3) and experimentally studied at length already (4–6), in the present work, we expand the scope of study to focus on subtle and as yet unexplored aspects of the interplay of biochemical noise -stochasticity- with time-scale separation between transcription and translation.

**Figure 1.**
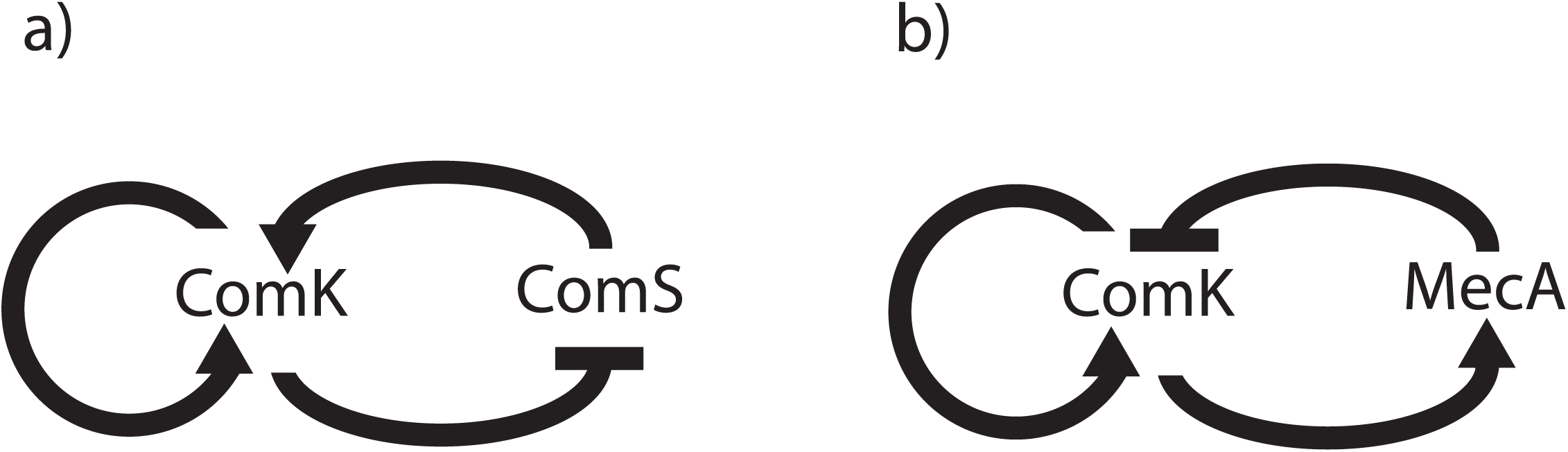
Core genetic regulation circuits. Panel a) depicts the competence phenotype core genetic regulation circuit of the canonical (native) *Bacillus subtilis* bacterium. Panel b) depicts a circuit with inverted indirect regulation. The term “inverted” refers to the inverted order of indirect feedback regulation compared to the canonical circuit. In the canonical circuit (panel a)), the indirect feedback regulation order is suppression followed by activation whereas in the inverted circuit (panel b)), the indirect feedback regulation order is opposite: activation followed by repression. ComK is the master regulatory gene of the competence phenotype; when ComK exceeds a certain threshold, the organism becomes competent to accept exogenous DNA into its own genome. In both the canonical and inverted circuits, ComK directly auto-activates itself. The organism controlled by the circuit depicted on panel a) is wild-type. The organism controlled by the circuit depicted on panel b), as far as we know, does not occur in Nature but it has been re-engineered in the laboratory (2). More details are in the text and references.

## Results

We conduct our investigations as much shepherded as bolstered by the concept demonstrated in (2) that dynamic oscillations are essentially competence phenotype breaking, and that the Evolution proofed core genetic regulation circuit of *Bacillus subtilis* must therefore operate in the excitatory regime. We thus consider these two dynamically close yet not biologically equivalent regimes, in the light of accessible evolutionary handles offered by stochasticity manifested as biochemical noise sourced in the paucity of some key molecular components in the system, and by time-scale separation between transcription and translation as it relates to effective time delays in the system.

### Canonical Bacillus subtilis regulation

Panel a) and panel b) of Figure 2 show the phase portraits of the “2D” infinite time-scale separation dynamics of the canonical *Bacillus subtilis* organism. The depicted dynamics is referred to as “2D” dynamics because, by stipulating the mRNA sub-manifold of a parent 4-dimensional system (ComK, ComS, mRNA_ComK_, mrRNA_ComS_) to be at rest (*i.e*. time derivatives of all mRNAs are all set to zero), the parent 4D system is collapsed into the two residual protein dimensions: ComK and ComS. Hence, mRNA does not appear in the dynamics in the 2D system. We refer to the time-scale separation (between transcription and translation) in the 2D system as “infinite”. This is the sense that increasingly slowing down mRNA translation effectively separates mRNA transcription from its translation by mimicking a time delay between the two processes. See Methods for more details.

**Figure 2.**
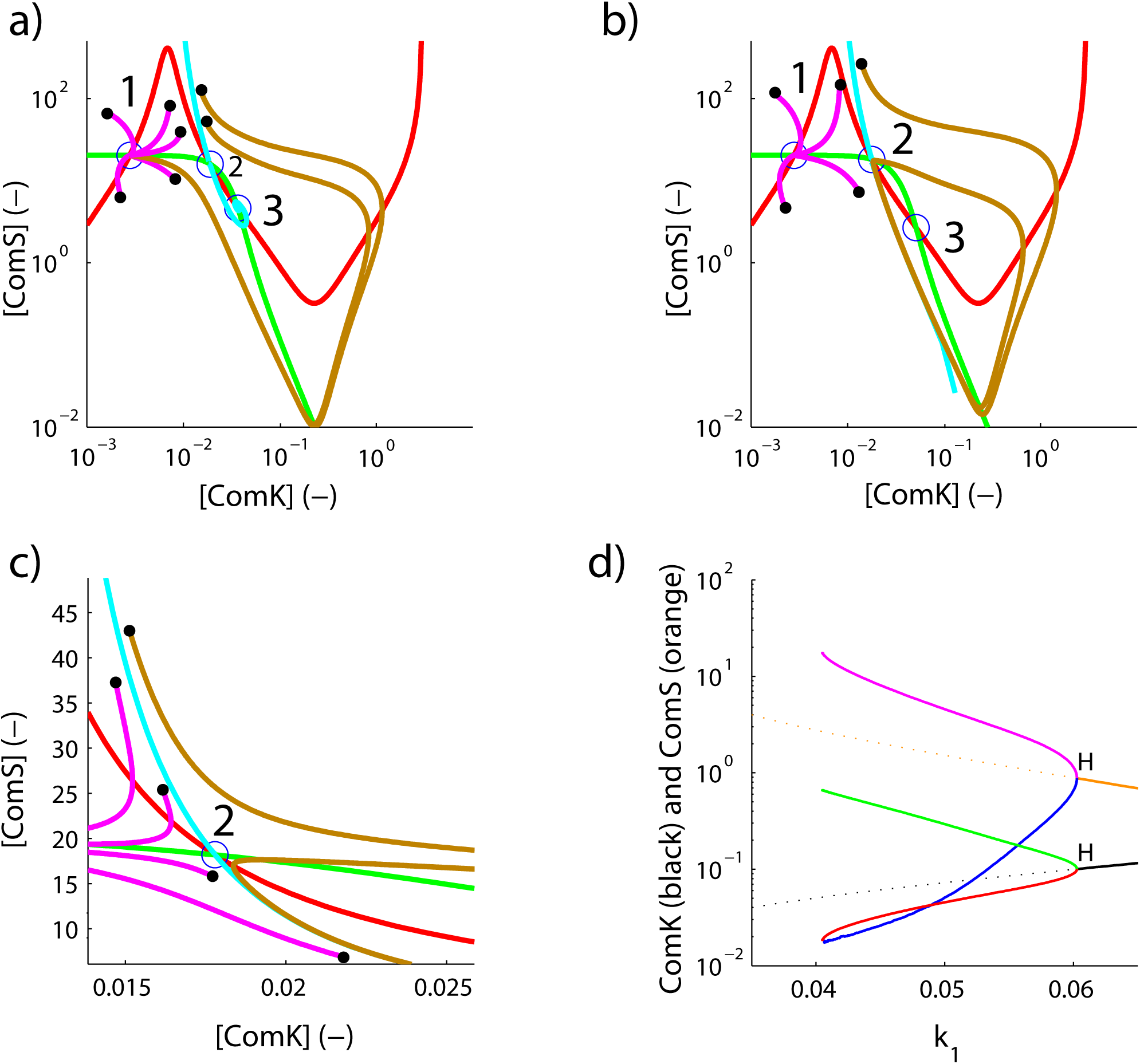
Canonical Bacillus Subtilis 2D circuit dynamics. Panels a) b) and c) are phase portraits with several sample tracks started at locations indicated by the black dots. The ComK nullcline is shown in red. The ComS nullcline is shown in green. Nullcline intersections indicated by the labels “1”, “2” and “3” are the fixed points of the dynamics. #1 is a stable spiral, #2 is a saddle node and #3 is an unstable spiral. On panel a), k_1_=.0333. On panel b), k_1_=.0405. On panels a) b) and c), the separatrix is shown by the cyan curve. Tracks (in magenta) started on the left of the separatrix on panels a) and b) fall into the stable fixed point #1. On panel a), tracks (in brown) started on the right side of the separatrix undergo an excitable trajectory about fixed point #3 and fall into fixed point #1. On panel b) however, similar tracks (in brown) also started on the right side of the separatrix, do not undergo an excitable trajectory. They instead fall into a limit cycle. Axes on panels a) and b) are in log base 10. Panel c) is a detailed view of panel b) in the region surrounding the saddle node #2. Axes on panel c) are linear. Panel d) shows the corresponding bifurcation diagram of the dynamics of fixed point #3 *vs*. k_1_. ComS is shown in orange. ComK is shown in black. Solid (dotted) orange and black lines indicate a stable (unstable) fixed point. A limit cycle clearly exists between the Hopf on the right (indicated by “H” at k_1_^∼^.06) and its appearance/disappearance point occurring on the left of the diagram (where the maximum amplitude purple and green curves and the minimum amplitude red and blue curves, all begin/end abruptly). At k_1_=.0333, there is no limit cycle (as on panel a). But at k_1_=.0405, the dynamics already exhibits a wide amplitude limit cycle, as on panel b) and panel c).

On Figure 2, panels a) b) and c), each diagram plots the genetic regulators ComK on the x-axis and ComS on the y-axis, and each plot features the two nullclines of the 2D dynamics (the ComK-nullcline, in red and the ComS nullcline in green). Nullclines are the loci of dynamical points on the phase diagram where the sub-manifold of ComK dynamics and the sub-manifold of ComS dynamics are individually, that is to say, separately, at rest. Therefore, at the intersections of nullclines, it is clear that the dynamics of the entire system is at rest. Here, the nullclines intersect at three fixed points: #1, a stable spiral, #2 a saddle node, and #3 an unstable spiral. Several tracks emanating at different initial conditions of ComK and ComS (black dots) illustrate the basic dynamical behaviors. We study the dynamics as a function of the key parameter of the feedback regulation, k_1_. The parameter k_1_ parameterizes the negative feedback of the ComK transcription factor onto the gene comS (see Methods). On panel a) at k_1_=.0333, magenta tracks fall into fixed point #1. Brown tracks started on the other side of the separatrix drawn in cyan, undergo a classic excitatory excursion around unstable fixed point #3 and eventually also reach fixed point #1. But on panel b) at) k_1_=.0405, the excitatory regime is replaced by a oscillatory regime; the brown track started on the right side of the separatrix now falls into a large amplitude limit cycle. On panel c), a blowup region of panel b), the details of the vicinity of the saddle node are shown in the case of the oscillatory track with k_1_=.0405. Whereas the excitatory biological regime of the canonical *Bacillus subtilis* is as on panel a), we notice that, actually, the excitatory regime lies not far at all from the oscillatory regime shown on panel b) and panel c). Indeed, regular large-amplitude oscillations suddenly replace excitatory trajectories as k_1_ increases from .0333 to .0405. It is clear that the implied phenotypes are drastically different between excitatory and oscillatory core genetic control. From the point of view of biology, the phenotype of interest is the appearance of core genetic regulation induced sustained yet transitory intervals of high levels of ComK. During high ComK levels, *Bacillus subtilis* is competent for accepting exogenous DNA (7–9). In the excitatory case, as depicted on panel a), random biochemical fluctuations must first occur in order to trigger the phenotype's randomly occurring and random-duration dynamical excursions into high ComK (brown tracks on panel a)). Indeed, it is easy to see that intrinsic randomness is required because if the system is started anywhere on the left side of the separatrix (cyan line), dynamics will only bring it to the lower fixed point #1, without excursions into the high ComK regime. Only upon a fluctuation that brings the starting point across to the right side of the separatrix, is the system then permitted to complete a large ComK excursion around fixed point #3. In the oscillatory case however, as shown on panel b) and panel c), high ComK intervals will also occur but at completely predictable and regularly spaced intervals of time. Whereas in the excitatory case, stochasticity is central to the dynamics, in the oscillatory case however, appears to be relegated to a blurring function.

No other tool than a fixed-point continuation analysis of the dynamics as shown on panel d) can summarize the behavior better. Here, k_1_ is the continuation parameter on the x-axis, and in black and orange are shown the locations and stability (solid: stable, dash: unstable) of the ComK and ComS upper fixed point #3. The values of ComK and ComS are jointly shown on the y-axis. There is a Hopf bifurcation denoted by “H” at k_1_∼.06. To the left of the Hopf bifurcation, oscillations are seen to develop. The upper/lower extrema of these oscillations are shown by the purple/blue and green/red lines for ComS and ComK, respectively. The amplitudes of these oscillations grow steadily from the Hopf as k_1_ diminishes until oscillations disappear abruptly towards the left of the figure. We refer to this point as the appearance/disappearance of oscillations because as k_1_ increases across this point, suddenly maximum amplitude oscillations appear, whereas as k_1_ is decreased across this point, the oscillations suddenly disappear. Oscillations occur due to the presence of the saddle, as detailed on panel c). To the left of the oscillation appearance/disappearance point (as on panel a)), the system is in excitatory mode. To the right (as on panel b) and panel c)), the system is in oscillatory mode. As stated above, to the left of this point, stochasticity is paramount to the canonical phenotype. To the right, is secondary.

### Looking for oscillations: the effects of time-scale separation and stochasticity

In this work, we are interested to study out how time-scale separation and stochasticity conspire to affect phenotype. We focus on the oscillatory regime because the phenotype is simple to spot, and appearance of oscillations in an otherwise excitatory regime represents competence-phenotype-breaking dynamical behavior and thus has a distinctly biological meaning. Figure 3 panel a) and panel b) show side by side a purely deterministic 2D (infinite time-scale separation) phase portrait and a discrete event stochastic simulation of imperfect (*i.e*. finite) time-scale separation. In both cases, k_1_=.05, a value that is expected to produce healthy oscillations of large amplitude. See Figure 2, panel c) for details. The limit-cycle on panel a) is unmistakable. On panel b), no limit cycle behavior can be seen. The stochastic dynamics is devoid of rotation; the system is still excitatory. We reasoned that the primary difference between these two regimes must be the specific amount of time-scale separation in the simulations shown on panel b); specifically that there might be insufficient timescale separation in the stochastic system.

**Figure 3.**
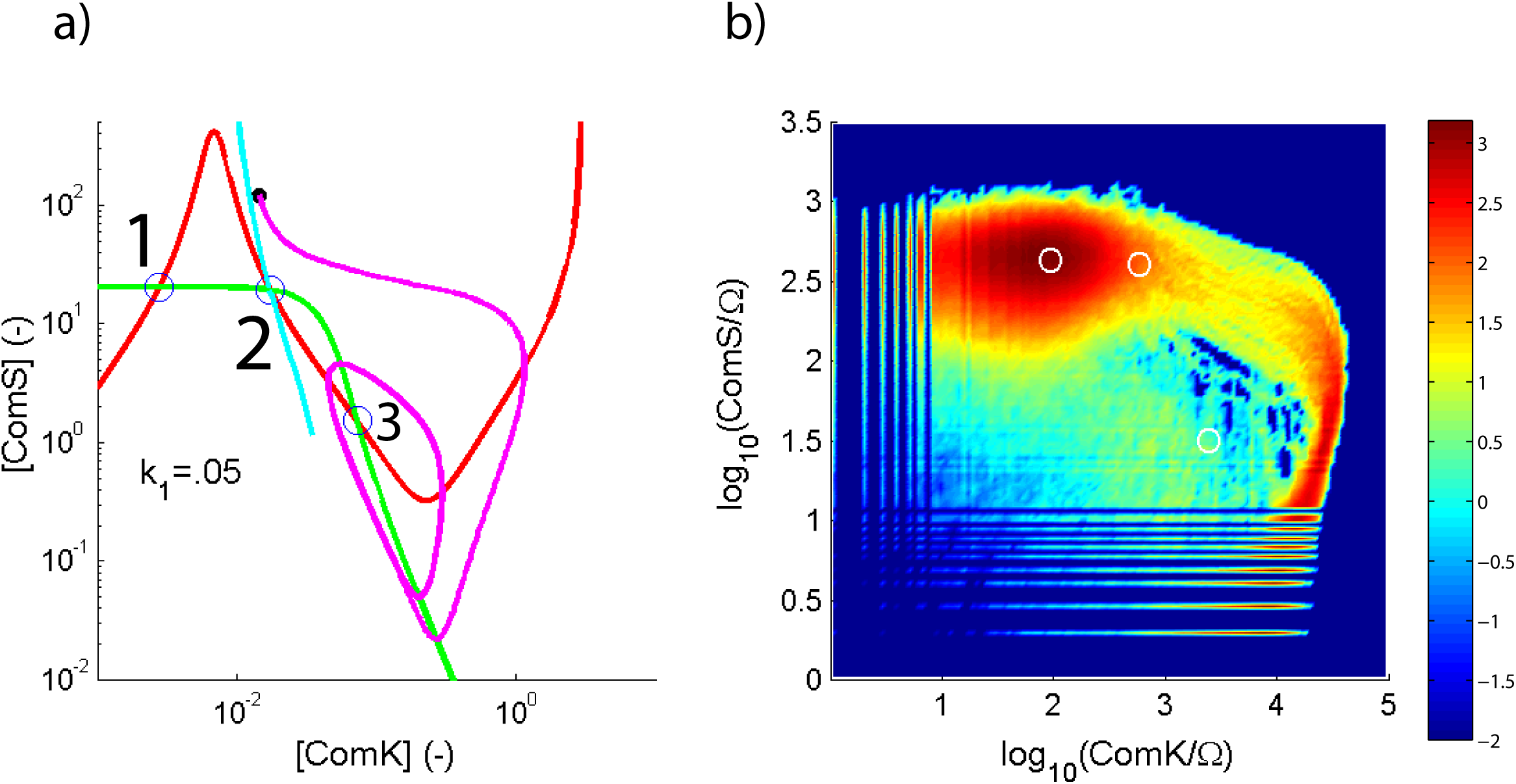
Deterministic and stochastic phase portraits of the canonical Bacillus subtilis circuit. Panel a), deterministic 2D phase portrait at k_1_=.05, well in the presence of the limit cycle (see panel c) of Figure 1). The ComK and Coms nullclines are the curves in red and green, respectively. Their intersections (“1”, “2” and “3”) are fixed points of the dynamics. Point #1 is a stable spiral, point #2 is a saddle node, and point #3 is unstable spiral. The magenta curve is a 2D computation of a trajectory started on the right side of the separatrix (shown in cyan), at the location indicated by the black dot. In this 2D simulation, by construction as explained in the text, the mRNA sub-manifold is at rest; time-scale separation is infinite. The limit cycle is obvious. The units on panel a) are dimensionless. Panel b) shows the corresponding stochastic simulation of the same system but, as explained in the text, by construction, the mRNA dynamics is not constrained to be at rest. In this particular case, time-scale separation is finite. Unlike panel a), the units on panel b) are dimensioned but for ease of comparison, the locations of the 2D fixed points (from panel a)) are shown on panel b) by three white circles. The color on the plot is a measure of the probability density for the stochastic system to occupy a certain state. The color bar on the right indicates the log base 10 value of the density corresponding to each color in the range. It is clear that the dynamics of the stochastic system, at this level of time-scale separation does not exhibit a limit cycle.

In order study the effect of time-scale separation unfettered by the possible influence of fluctuations, we devised a fluctuation-free version of the stochastic system that we refer to as the “6D system”. This continuous dynamical system is derived from the set of discrete-event reaction equations underlying the stochastic system, but completely removing the fluctuations. In effect, the 6D system is the infinite number of molecule limit (i.e. so-called thermodynamic limit) of the stochastic system; hence 6D is a deterministic system. More details are given in the Methods section. We aimed to study the impact of time-scale separation on the dynamics and to do so we conducted bifurcation studies versus translation slowdown factors (SDT_ComK_ and SDT_ComS_) introduced in the dynamics. More details are in Methods. Figure 4 show the 6D system's fixed point locations as we vary the translation factors slowdown. The presence of the Hopf bifurcation testifies that oscillations are clearly induced by slowing down the mRNA translation sub-manifold. We picked SDT_ComS_ = .9 and SDT_ComK_ = .14 from these studies because these factors correspond to steady mid-range healthy oscillations well away from critical points.

**Figure 4.**
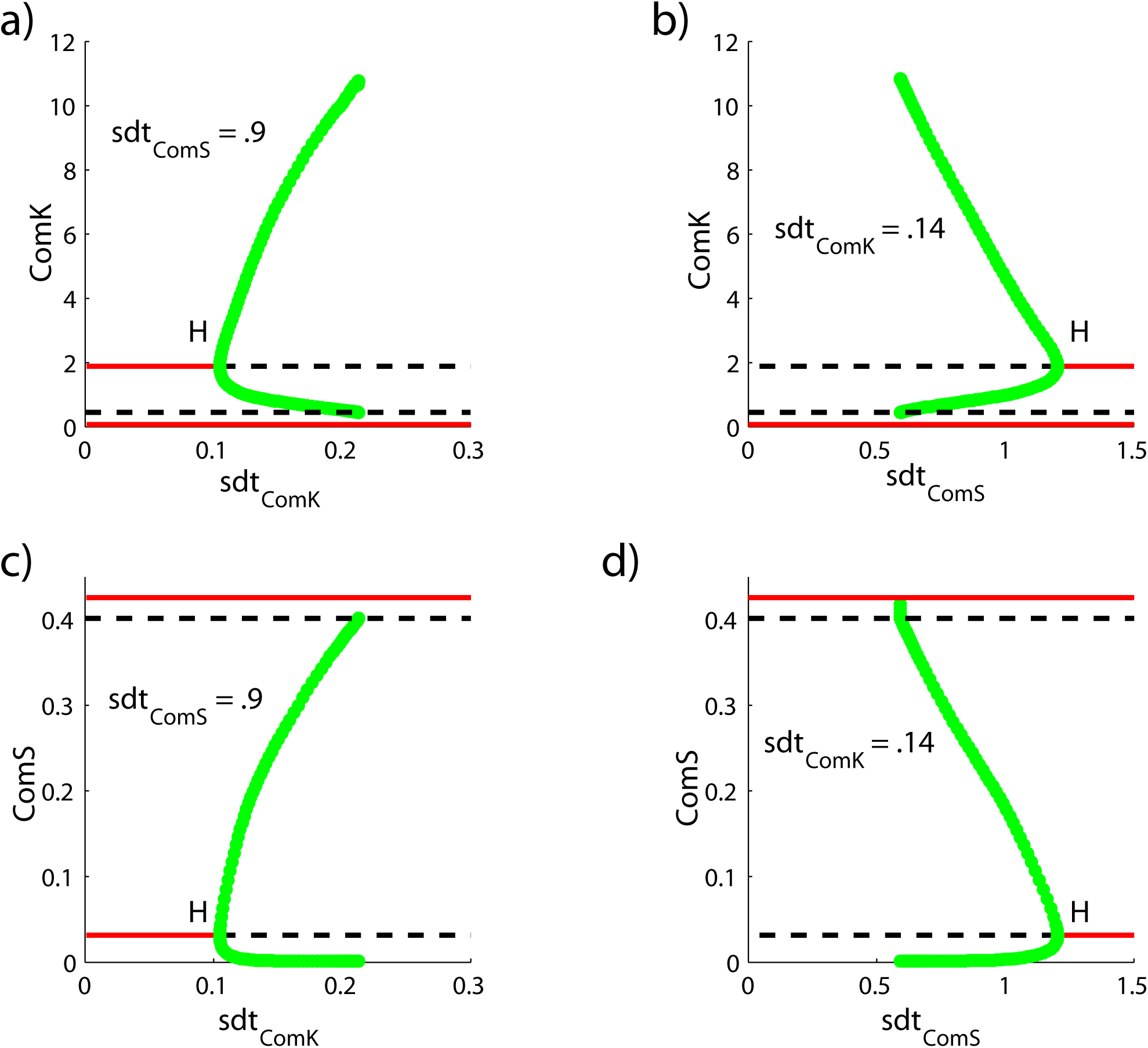
Dynamical behavior of 6D system vs. slowing down of translation (SDT) factors. As explained in the text, in order to facilitate dynamical exploration, a 6D deterministic system was derived from the stochastic discrete event system. This cognate 6D system corresponds to the thermodynamic limit of the stochastic system: it is the infinite number of molecule limit of the stochastic system. The 6D system displays the evolution of the mean of the stochastic system in the limit of infinite number of molecules, therefore with zero fluctuations. Panel a) and b) are bifurcation diagrams of the 6D system showing the ComK fixed point locations *vs*. two slowing down factors on the translation manifolds: SDT_ComK_ acting on the ComK mRNA sub-manifold and SDT_ComS_ acting on the ComS sub-manifold. On panel a), SDT_ComS_ = 0.9, and on panel b), SDT_ComK_ = 0.14. Panel c) and panel d) are the equivalent bifurcation diagrams for ComS. All bifurcation diagrams exhibit a Hopf bifurcation indicated by “H” and delineate the dynamical regions where rotation exists (low and high amplitudes of the concomitant limit cycle are shown in green). These four diagrams make it emphatically clear that the choice of SDT_ComS_ = 0.9 and SDT_ComK_ = 0.14 are slowing down translation factors on the dynamics of the 6D system that result in healthy limit cycle dynamics located in mid-range of amplitude, away from the Hopf and away from the appearance/disappearance points of oscillations (end points of green curves on right side of the diagram). The diagrams show stable fixed points in solid red or black, and unstable fixed points in dashed red or black. The fixed point with the Hopf bifurcation and associated rotation are denoted by #3 on Figure 2 and on Figure 3.

On Figure 5, panel a) we show, in magenta, a 2D track (with infinite time-scale separation) falling into the expected limit cycle, and we also show (in cyan) a 6D track with SDT_ComS_ = .9 and SDT_ComK_ = .14 resulting in similar oscillations. Thus the 2D system and the 6D system are in agreement. Note that a 6D track without translation slowdown (*i.e*. with translation slow down factors SDT_ComS_ and SDT_ComK_ set to 1) fails to fall into oscillations, as is demonstrated on Appendix Figure 1. Panel b) shows the equivalent stochastic computation with the slow down factors applied. Despite the presence of translation slowdowns in the stochastic simulation, it is abundantly clear that no oscillations are induced yet; time-scale separation that brings the dynamics to oscillations in the deterministic infinite-number of molecule limit represented by the 6D system is clearly insufficient to bring the corresponding stochastic system with finite stochasticity into similar oscillations. Clearly stochasticity level, being the other parameter in the system, must be impacting the phenotype. This is shown by the simulation presented on panel c). Here, as on panel b) slowdown of translation is in effect, but in addition, the system×s intrinsic stochasticity level is lowered by roughly a factor of 3 compared to panel b). We observe the clear unmistakable occurrence of oscillations.

**Figure 5.**
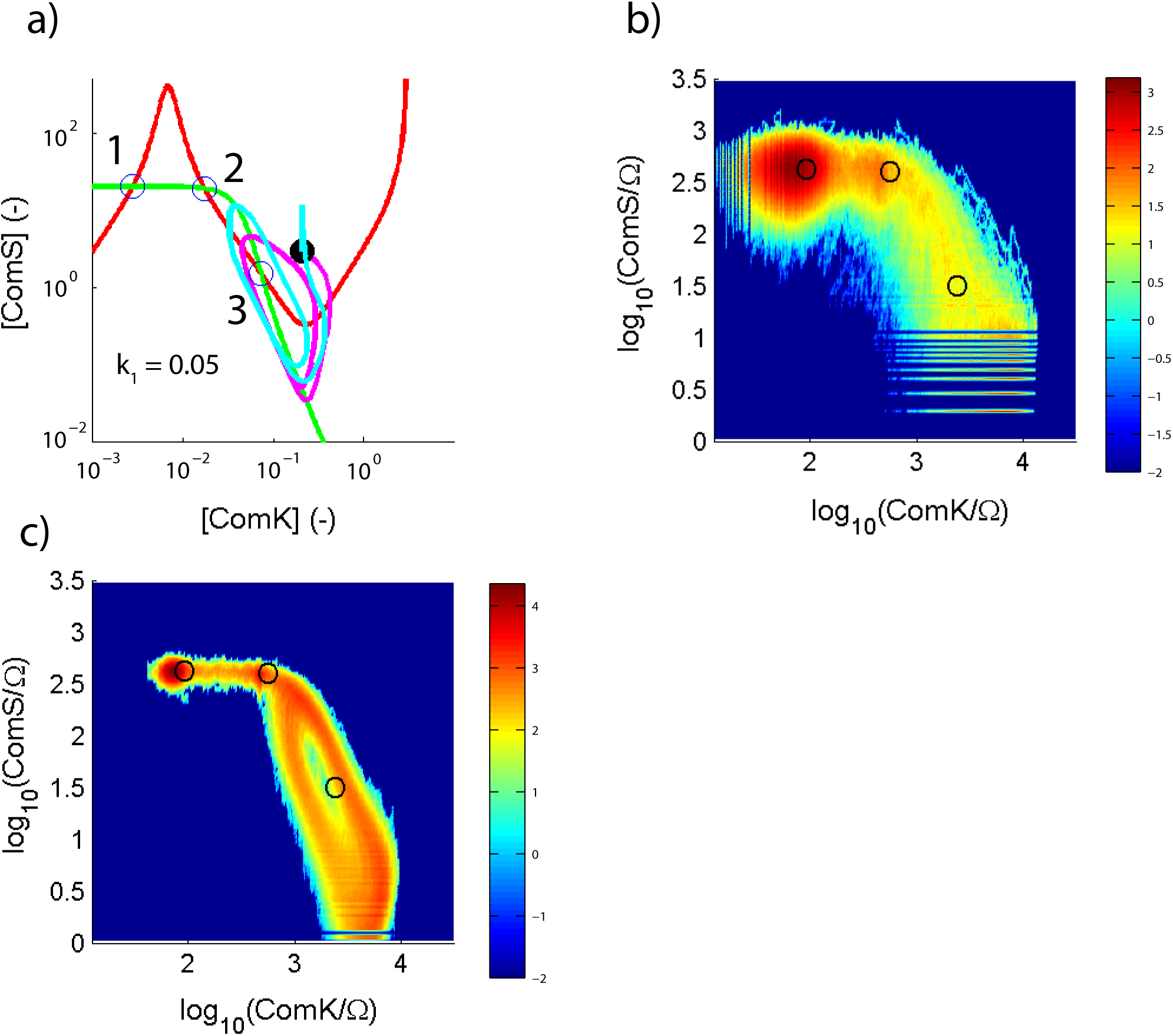
Effect of increasing time-scale separation by slowing down translation (SDT) and effect of reducing fluctuations. Panel a) shows the phase portrait of the canonical Bacillus subtilis circuit with k_1_ = .05. The nullclines and the fixed points of the dynamics are the same as on Figure 3, panel a). Here, a 2D computation of a trajectory initiated at the location of the black dot is shown in magenta. The 2D system exhibits built-in infinite time-scale separation: the mRNA sub-manifold is at rest. The 2D magenta track clearly falls into a limit cycle. Shown in cyan is the corresponding 6D computation (also initiated at the same location as the 2D) for which the slowdown factors SDT_ComS_ = 0.9 and SDT_ComK_ = 0.14 are enforced. The values for the slowdown factors are determined using bifurcation analysis (see Figure 4). As explained in the text, the 6D system is the infinite number of molecule limit of the corresponding stochastic system, shown on panel b). In the 6D system, while the mRNA sub-manifold is not at rest by construction, the presence of the slowdown factors on the translation dynamics enforces a large time-scale separation that brings the behavior close to that of the 2D system. Therefore, the 6D endowed with translation slowdown cyan track also falls into a limit cycle. Note that a 6D track without translation slowdown (SDT_ComS_ = 1 and SDT_ComK_ = 1) is excitatory (see Figure AF3). As stated above, panel b) shows the corresponding stochastic simulation with the same translation slowdowns as on panel a) but with a canonical level of noise. It is clear that, in this noise regime, there is no limit cycle. Panel c) however shows the stochastic simulation with the translation slowdown factors but for which both the number of molecules and the simulation volume are multiplied by a factor of 10, thus leaving the concentrations intact yet reducing the intrinsic fluctuations by roughly 10^1/2^. As expected, the limit cycle appears. Thus both time-scale separation and biochemical noise conspire to hide oscillatory dynamics. Panel a) is in dimensionless unit and panel b) and panel c) are in dimensioned units. For convenience, on panel b) and panel c), the locations of the 2D system dynamical fixed points from panel a) are indicated by black circles.

**Figure AF1.**
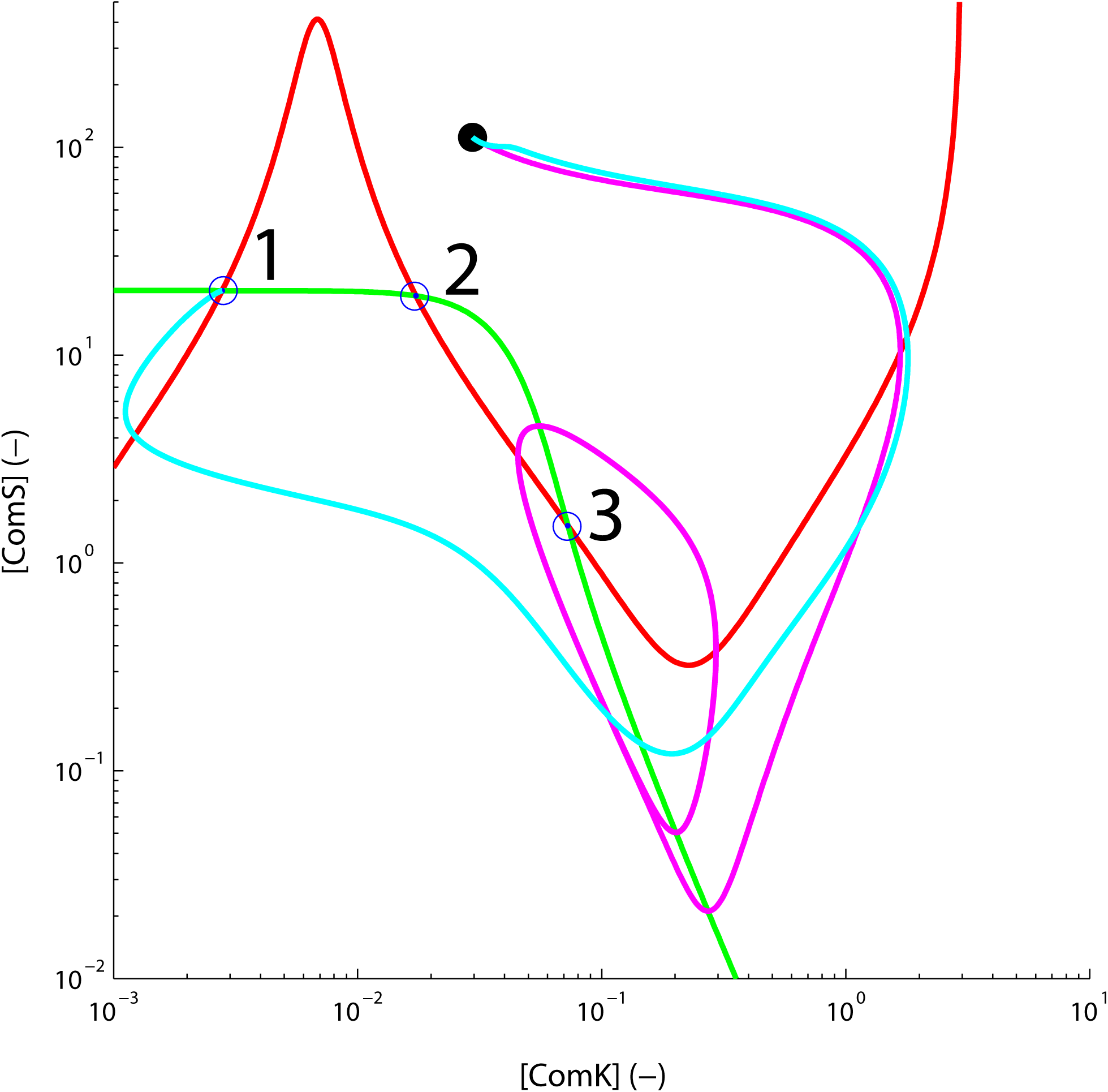
2D computation and 6D computation without translation slowdowns. Phase diagram of the canonical Bacillus subtilis system with k_1_ = .05. The ComK and Coms nullclines are the curves in red and green, respectively. Their intersections (“1”, “2” and “3”) are fixed points of the dynamics. Point #1 is a stable spiral, point #2 is a saddle node, and point #3 is unstable spiral. The magenta curve is a 2D computation for which the mRNA sub-manifold is at rest and therefore, time-separation is infinite. The dynamics is oscillatory; the magenta track originating at the location of the black dot falls into a limit cycle. The cyan line is a 6D computation devoid of translation slowdown (SDT_ComS_ = 1 and SDT_ComK_ = 1). The 6D system displays the evolution of the mean of the corresponding stochastic system in the limit of infinite number of molecules, therefore with zero fluctuations. The 6D dynamics is excitatory; the cyan track also originating at the location of the black dot does not fall into a limit cycle, but instead heads back to the stable fixed point #1.

Thus both time-scale separation and stochasticity conspire to mask the oscillation phenotype; or, from a biological viewpoint, time-scale separation and stochasticity work together to maintain the system in an excitatory regime exhibiting the competence phenotype.

### Inverted *Bacillus subtilis* regulation

Hereon, we shift focus to the inverted *Bacillus subtilis* regulation. This circuit was not selected by Evolution for reasons explained in (2). However, because it was genetically re-engineered for investigation, the experimental behavior of the inverted *Bacillus subtilis* circuit is well understood. Figure 6, panel a) and panel b) present the phase portrait of the 2D (parent 4D system's mRNA sub-manifold at rest) regulation in which the two regulators are ComK and MecA. Nullclines for these two regulators are shown in red and green, respectively. The three fixed points of the dynamics, located at their mutual intersections, are shown by the labels “1”, “2” and “3”. They are stable spiral, saddle point, and unstable spiral, respectively. The separatrix is shown in cyan. On panel a), k_m_=.1474 (excitatory regime, no limit cycle) and on panel b), k_m_=.1475 (oscillatory regime, large limit cycle). The parameter k_m_ is key to the feedback regulation as it parametrizes the positive feedback of transcription factor ComK onto the gene mecA (see Methods). Computed trajectories are shown in magenta. They originate at the location of the black dot. Note that the direction of motion (here, counterclockwise) is opposite that of the canonical circuit (clockwise, see Figure 2, for example). Figure 6, panel c) shows the bifurcation analysis of the 2D system vs. k_m_. It depicts the overall dynamics and, in particular, it explains the sudden appearance of the large limit cycle observed between k_m_=.1474 and k_m_=.1475. There is a Hopf bifurcation denoted by “H” at k_m_ ∼.1559 where oscillations are born. The ComK and MecA locations of fixed point #3 are indicated by black and orange lines, respectively. Solid lines are used to indicate stability while dotted lines are for unstable. As k_m_ decreases, the amplitude of the limit cycle born at the Hopf increases as is denoted by showing the limit cycle's upper and lower limits: purple and blue lines for MecA; green and orange lines for ComK. On the left side of the diagram, these oscillations terminate abruptly. Again, as in the case of the canonical *Bacillus subtilis* regulation circuit, we refer to this point at the oscillation appearance/disappearance point depending on whether crossing is from the left to right (increasing k_m_) or opposite direction. In fact these oscillations appear in a similar dynamical manner as for the canonical circuit, that is to say, they appear at the saddle node, hence similar to Figure 2c). However, the detailed view for the inverted circuit is omitted for brevity.

**Figure 6.**
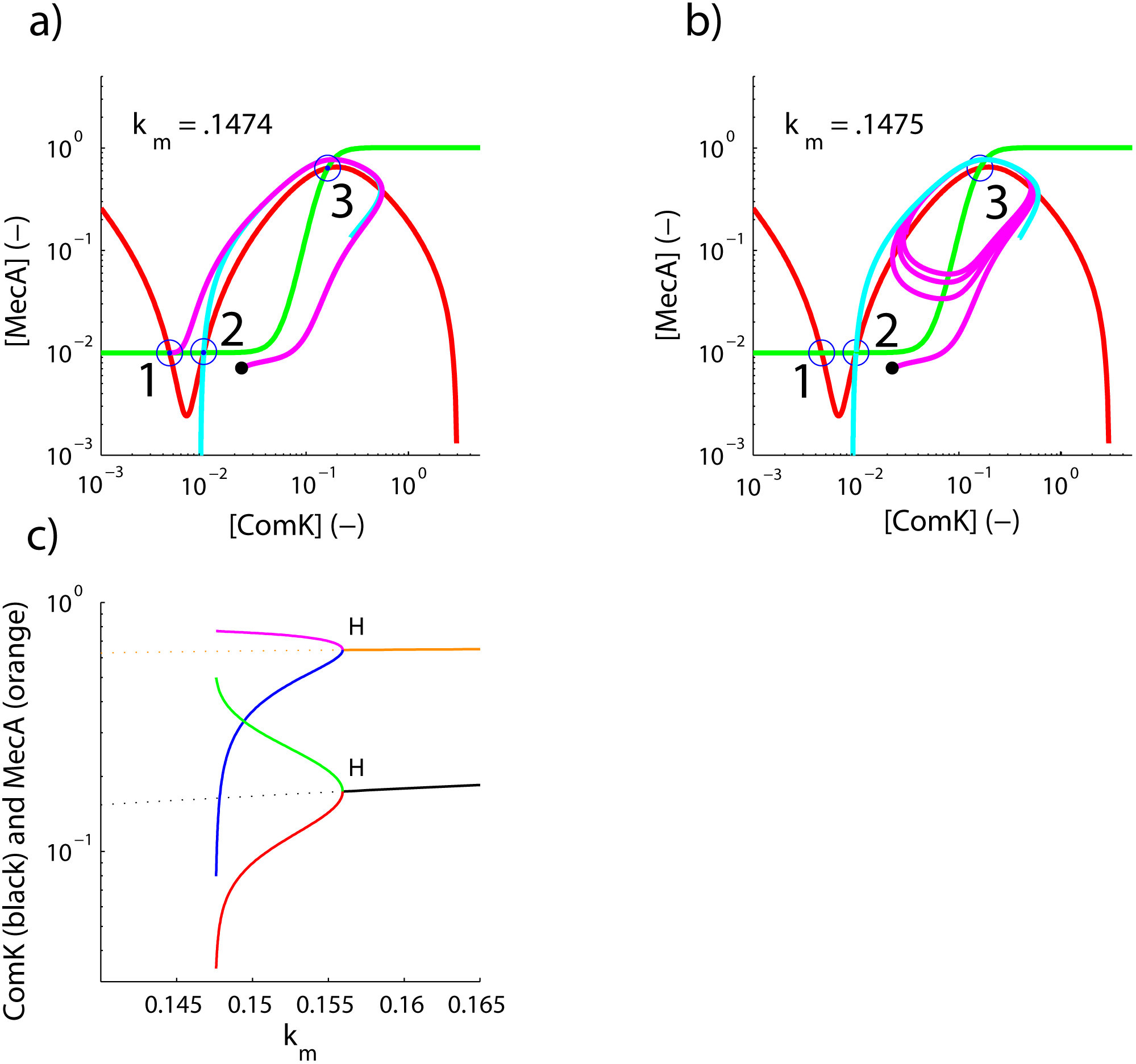
2D Dynamics of the inverted Bacillus subtilis circuit. The ComK nullcline is shown in red and the MecA nullcline is shown in green. The nullcline intersections labeled “1”, “2” and “3” are stable spiral, saddle, and unstable spiral fixed points of the dynamics, respectively. The location of the separatrix is indicated by the cyan curve. On panel a), k_m_ = .1474 and on panel b) k_m_ = .1475. On panel a) and b) a 2D system track is started at the same location indicated by a black dot. On panel a) the system is the excitatory regime so the track winds around fixed point #3 and falls back into fixed point #1. On panel b) however, the system is in oscillatory mode so the track falls into a limit cycle. Panel a) and b) have dimensionless units. Panel c) is a bifurcation analysis of the 2D dynamics showing in orange and red the location of fixed point #3 for MecA and ComK, respectively. Solid lines indicate this fixed point is stable; dashed lines indicate the fixed point is unstable. There is a Hopf bifurcation located at k_m_ ^∼^ .1559. To the left of the Hofp develops a limit cycle whose maximum/minimum amplitudes are denoted in purple/blue and green/red lines for MecA and ComK, respectively. Towards the left of the diagram is the appearance/disappearance point of the limit cycle located between k_m_ = .1474 and k_m_ = .1475. The dynamical behavior observed on panel a) and panel b) is therefore corroborated by the bifurcation analysis. At k_m_ = .1474 no limit cycle is expected while at k_m_ = .1475 one observes a large amplitude limit cycle. The 2D system has built-in infinite time-scale separation; its mRNA sub-manifold is at rest.

Figure 7, panel a), panel b) and panel c) show corresponding stochastic simulations at k_m_= .1, k_m_ = 1.5 and k_m_ = .2 respectively. Note that since units on the stochastic simulation plots are dimensioned, a scaling factor “sf” appears such that the effective K_m_ = k_m_ X sf (see Methods). Appendix Figure 2 details how these probability density plots are obtained from combining a large number of statistically independent histories (see Methods). In accordance to the 2D bifurcation analysis shown on Figure 6, panel c), we observe the stochastic simulations shown on panel a), panel b) and panel c) of Figure 7 to be in excitatory, oscillatory and bi-stable regimes, respectively. Thus, there is general dynamical agreement between the two viewpoints: deterministic and stochastic. However, the deterministic dynamics was computed, by design, in the limit of the mRNA submanifold being at rest (infinite time-scale separation) whereas, the stochastic regimes shown on Figure 7 do not strictly enforce this limit. On Figure 7, the stochastic simulations have finite time-scale separation, not infinite. Below we focus on some detailed consequences of this difference: the two viewpoints (deterministic and stochastic) do, in fact, lack specific dynamical agreement.

**Figure 7.**
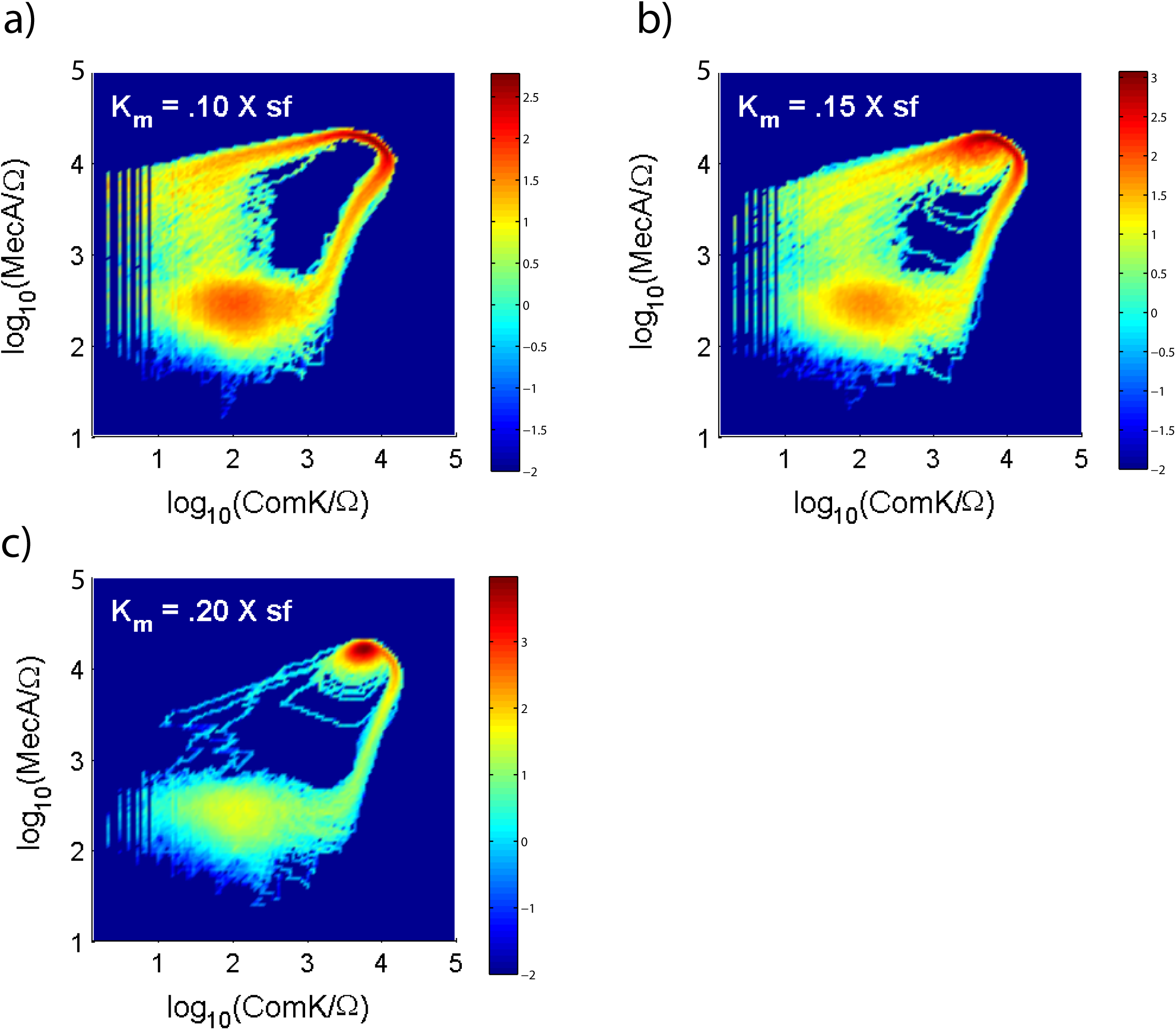
Stochastic dynamics of the inverted Bacillus subtilis circuit. The three panels show probability densities as a function of the k_m_. In panel a), k_m_ = .1, on panel b), k_m_=.15 and on panel c) k_m_= .2. The units on these plots are dimensioned so the effective K_M_ = k_m_ × sf where the scaling factor “sf” takes care of dimensions. On panel a) the system is in excitatory regime, on panel b) the system is in oscillatory regime (notice the presence of limit cycle behavior) and on panel c) the system is bi-stable as the upper fixed point's stability has now shifted to stable spiral. These three dynamical regimes are roughly predicted by the bifurcation analysis shown on Figure 6, panel c). However, the correspondence is only expected to be approximate because whereas in the 2D system the time-scale separation is infinite, in the canonical system shown here, it is finite. The colorbars on the right of each plot indicate the log base 10 of the density.

**Figure AF2.**
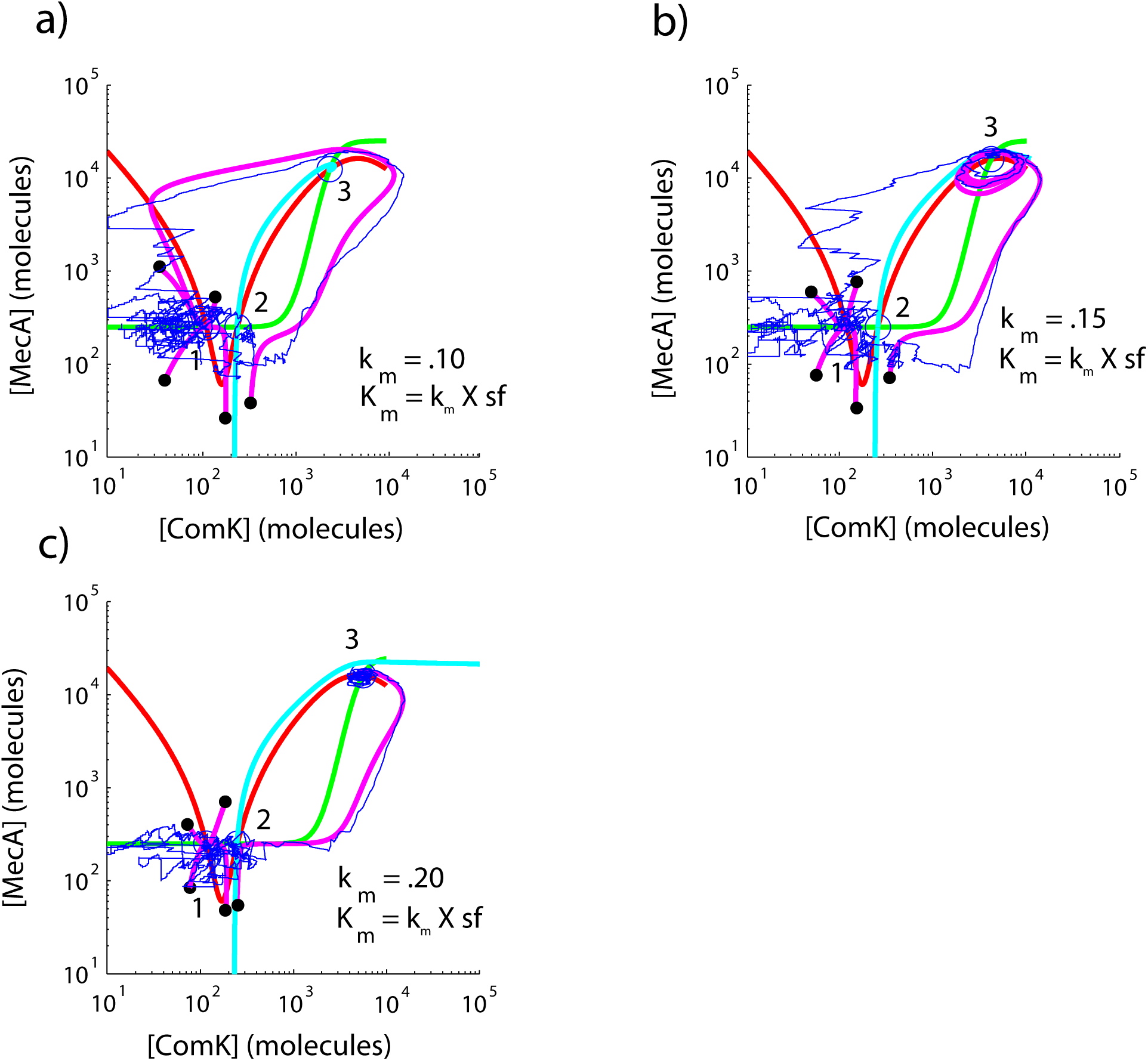
Inverted Bacillus subtilis deterministic and stochastic dynamics. The ComK nullcline is shown in red and the MecA nullcline is shown in green. The nullcline intersections labeled “1”, “2” and “3” are stable spiral, saddle, and unstable (panel a and b)/stable (panel c) spiral fixed points of the dynamics, respectively. The location of the separatrix is indicated by the cyan curve. On panel a), k_m_ = .1, on panel b) k_m_ = .15 and on panel c) k_m_ = .2. The units on these plots are molecules in a standard unit volume. The stated k_m_ can be used to locate the expected dynamical regime from the bifurcation analysis shown on Figure 6, panel c). On panel a), the 2D dynamics is excitatory, on panel b) the 2D dynamics is oscillatory and on panel c), passed the Hopf bifurcation, the 2D dynamics is bi-stable (the upper fixed point #3 is now stable). The purple tracks on these plots are several 2D computations initiated at the same locations on each diagram indicated by the black dots. Each blue track on panel a), panel b) and panel c) is a single instance of a stochastic track for which the intrinsic time-scale separation, unlike the 2D deterministic system, is not infinite. Nevertheless, in these particular regimes, there is good qualitative agreement between the 2D deterministic and stochastic computations. The starting point of the stochastic track is the same on all panels and is located in the general vicinity of fixed point #1. The track first meanders in the vicinity of fixed point #1 for some time in all regimes. On panel a) the track then undergoes a classic excitatory excursion. On panel b), the track gets trapped momentarily in the limit cycle where it undergoes several oscillations, but it eventually escapes the limit cycle. In the regimes depicted on panel a) and b), the track eventually comes back to the vicinity of fixed point #1. However, on panel c), the track gets permanently trapped in the now stable upper fixed point #3. Probability densities (occupancy diagrams) are obtained from adding together the effect of 200 statistically independent extended stochastic simulations.

### Persisting oscillations in finite time-scale separation

We may ask how the finite time-scale separation affects the stochastic dynamics by focusing on the oscillation appearance/disappearance point (left side of Figure 6, panel c)). For reference, on Figure 8 panel a) we show the finite time-scale separation stochastic simulation at k_m_ = .125, that is to say, in robust excitatory regime (refer to Figure 6, panel c)). There is no limit cycle in the dynamics. On panel b) and panel c), k_m_ = .1474 and k_m_ = .1475, respectively. From the 2D bifurcation analysis, at k_m_ = .1474 there should be no oscillations, and at k_m_ = .1475 large amplitude oscillations are expected. However, panel b) and panel c) of the stochastic dynamics actually depict similar oscillatory behavior at these two values of k_m_: the clear oscillations around the upper fixed point seen on panel c) also persist to the lower k_m_ regime of panel b). We seek the source of the discrepancy.

**Figure 8.**
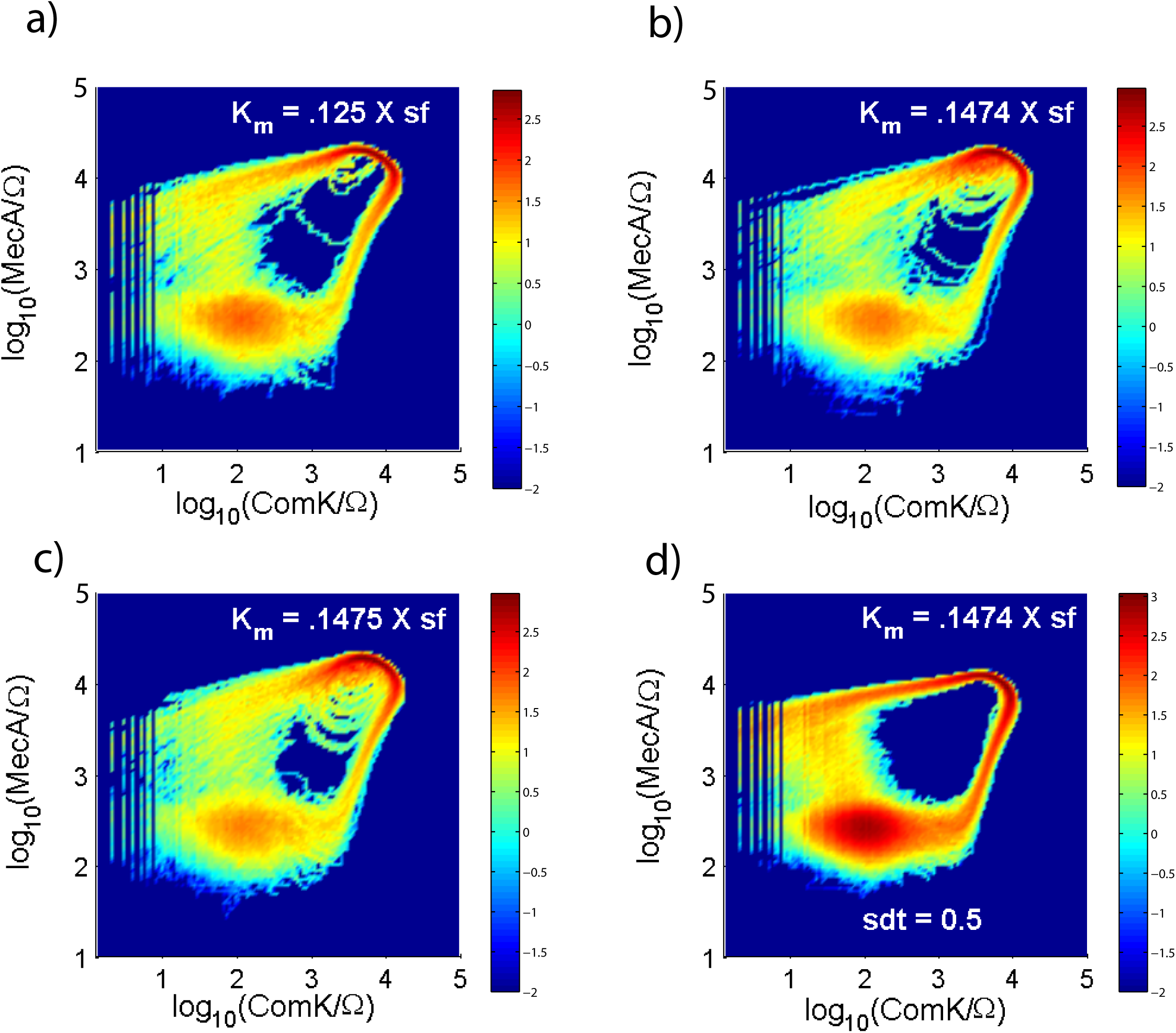
Appearance/disappearance of the limit cycle in the Stochastic dynamics of the inverted Bacillus subtilis circuit. The three panels show probability densities as a function of the k_m_. In panel a), k_m_ = .125, on panel b), k_m_=.1474 and on panel c) k_m_= .1475. The units on these plots are dimensioned so the effective K_M_ = k_m_ × sf where the scaling factor “sf” takes care of dimensions. On panel a), as expected from the 2D bifurcation analysis (Figure 6, panel c)), the dynamics does not exhibit a limit cycle: it is purely excitatory. On panel b), just before the appearance/disappearance point of the limit cycle, and on panel c) just following it, both diagrams exhibit similar limit cycle behavior. Whereas on panel b), according to the 2D bifurcation analysis, there should not be any limit cycle behavior, there clearly is, as much as there is on panel c) where oscillations are expected and observed. Panel d) offers a stochastic simulation also for k_m_=.1474 but including a translation slowdown factor SDT=0.5. Comparing panel b) to panel d), it is clear that the inclusion, on panel d), of significant time-scale separation in the system makes the limit cycle behavior disappear. Hence, the stochastic system with translation slowdown has acquired increased similarity with the 2D infinite time-scale simulation.

Focusing on the time-scale separation difference between the 2D computation and the stochastic simulation, we show on panel d) of Figure 8 a stochastic simulation that now includes a translation slowdown factor SDT = 0.5 (common to both ComK and MecA) (see Methods). Comparing panel b) and d), it is emphatically clear that increasing time-scale separation makes the limit cycle disappear from the stochastic simulation: there is no cycling about the upper fixed point anymore. This behavior is as expected as the stochastic simulation's timescale separation is now closer to that of the 2D ideal. Could stochasticity alone account for the difference? Appendix Figure 3 shows a stochastic simulation without translation slowdown, but for which the number of molecules and volume are simultaneously increased by a factor of 10 thus leaving concentrations unchanged but reducing intrinsic fluctuations by about a factor of 3. A factor of 10 in number of molecules is arguably biologically maximal. Comparing Appendix Figure 3 panels a), b) and c) to the corresponding ones of Figure 8, we conclude that reduced biochemical noise alone cannot be the source of disagreement for the location of the oscillation appearance/disappearance point in the stochastic simulations. In fact, as we have shown above, it is time-scale separation that constitutes the source of disagreement (again, compare Figure 8, panel b) to panel d)). Hence, in the inverted *Bacillus subtilis* system, finite time scale separation extends the oscillatory regime range, or, from a biological point of view, finite time scale separation reduces the range of excitability.

**Figure AF3.**
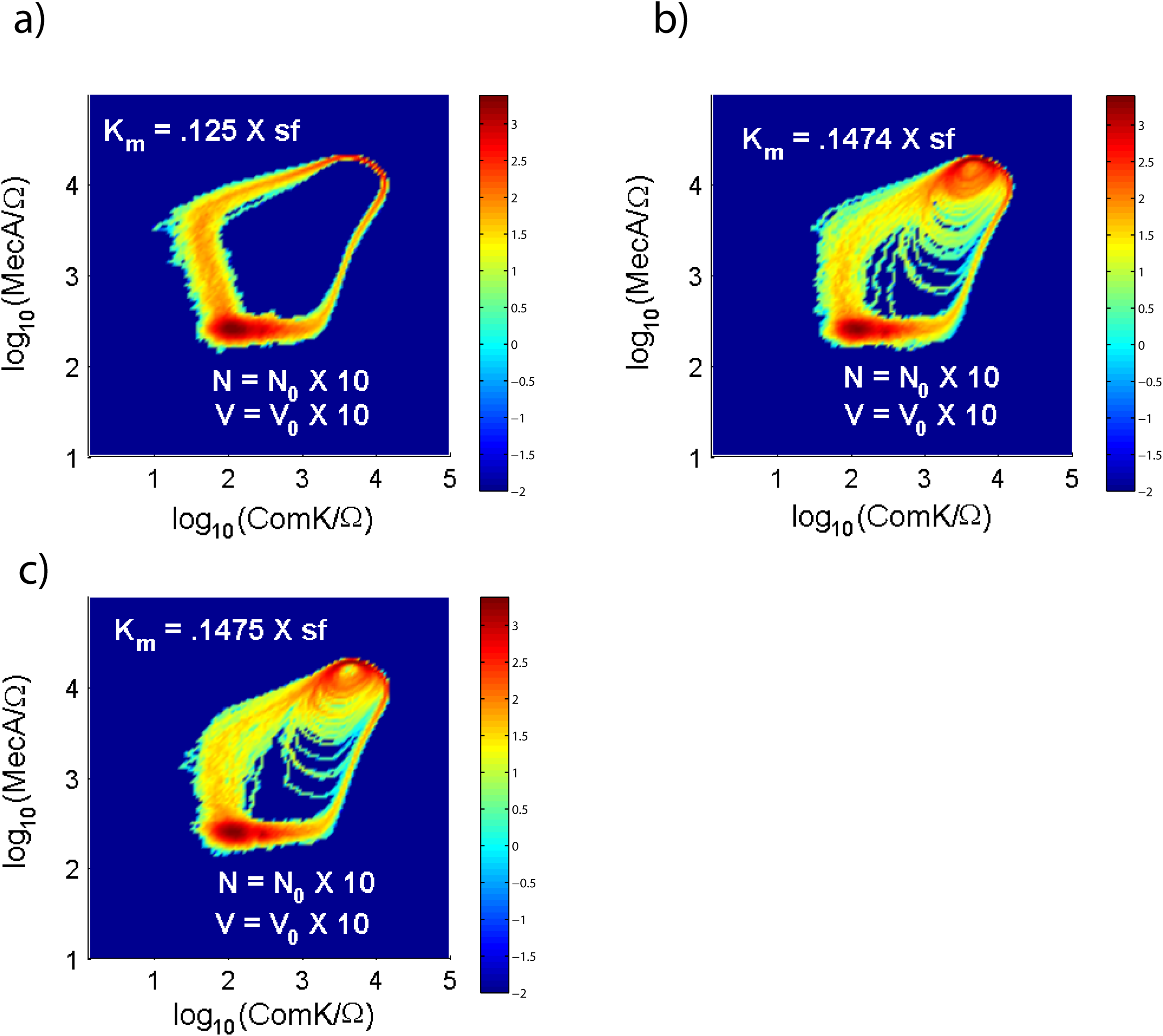
Less stochasticity does not induce appearance/disappearance of the limit cycle in the Stochastic dynamics of the inverted Bacillus subtilis circuit. The three panels show probability densities as a function of the k_m_. In panel a), k_m_ = .125, on panel b), k_m_=.1474 and on panel c) k_m_= .1475. The units on these plots are dimensioned so the effective K_M_ = k_m_ × sf where the scaling factor “sf” takes care of dimensions. Unlike Figure 8, all simulations are done with 10 times the number of molecules in 10 times the volume, such that molecular concentrations are left intact but stochastic variations are reduced by roughly 10^0.5^. On panel a), as expected from the 2D bifurcation analysis (Figure 6, panel c)), the dynamics does not exhibit a limit cycle: it is purely excitatory. On panel b), just before the appearance/disappearance point of the limit cycle, and on panel c) just following it, both diagrams exhibit similar limit cycle behavior. Whereas on panel b), according to the 2D bifurcation analysis, there should not be any limit cycle behavior, there clearly is, just as much as there is on panel c) where oscillations are expected and observed. Thus the clash between the 2D infinite time-scale separation computation and the stochastic simulation is not resolved by reducing the noise.

## Methods

### Canonical and inverted *Bacillus subtilis* circuits and the feedback continuation parameters

The deterministic and stochastic models of the canonical and inverted *Bacillus subtilis* circuits are thoroughly developed in (2). The canonical (*i.e*. native) *Bacillus subtilis* core circuit was devised to explain the “Meks” bacterium behavior whereas the inverted *Bacillus subtilis* circuit was devised to shed light on the “Synex” mutant behavior. In (6), the deterministic and stochastic models of the canonical *Bacillus subtilis* circuits are also presented in details. In this work we reuse exactly the same models; equations and parameters.

In the specific cases of the continuation parameter k_1_ of the deterministic canonical model and the corresponding parameter k_m_ of the deterministic inverted model, we varied these parameters over a range to produce the relevant bifurcation analyses. More specifically, the role of k_1_ is given by equation S3 in the supplement of (6), where it parameterizes the negative feedback onto the gene comS by the protein ComK. The role of k_m_ is similarly described by equation (2) of the supplement of (2). Here the regulation is that of positive feedback onto the gene mecA by the protein ComK.

### 6D model: the infinite number of molecule limit of the underlying stochastic model of the canonical *Bacillus subtilis* circuit

The equations for the 6D model are straightforwardly assembled, term by term (production terms are positively signed, degradation terms are negatively signed) from the set of discrete-event reactions of (2) and are given below.

~~~
d mRNA_ComK_ / dt = Pconstitutive_ComK_ x k_1_ + P_ComK_ x f(ComK,k_2_,k_k_,n) - mRNA_ComK_ x k_7_
~~~

~~~
d mRNA_ComS_ / dt = Pconstitutive_ComS_ x k_4_ + P_ComS_ x g(ComK,k_5_,k_s_,p) - mRNA_ComS_ x k_9_
~~~

~~~
d ComK / dt = mRNA_ComK_ x k_3_ + MecA_ComK x km_11_ - ComK x k_8_ - MecA_free_ x ComK x k_11_
~~~

~~~
d ComS / dt = mRNA_ComS_ x k_6_ + MecA_ComS x km_13_ - ComS x k_10_ - MecA_free_ x ComS x k_13_
~~~

~~~
d MecA_ComK / dt = MecA_free_ x ComK x k_11_ - MecA_ComK x km_11_ - MecA_ComK x k_12_
~~~

~~~
d MecA_ComS / dt = MecA_free_ x ComS x k_13_ - MecA_ComS x km_13_ - MecA_ComS x k_14_
~~~

in which

~~~
f(ComK,k_2_,k_k_,n) = (k_2_ x ComK^n^) / (k_k_^n^ + ComK^n^)
~~~

~~~
g(ComK,k_5_,k_s_,p) = k_5_ / (1 + (ComK/k_s_)^p^)
~~~

~~~
MecA_free_ = (M_T_ - MecA_ComK - MecA_ComS)
~~~

Because these equations have no innate fluctuations, they represent the infinite number of molecules version of the underlying stochastic model. They are expected to reproduce the time evolution of the first moment of the stochastic model state variable distributions that would be obtained with an infinite number of statistically independent histories. The parameters of the 6D model are consistent with the stochastic model in [(2). For convenience, we reproduce them below; in dimensioned units:

~~~
k_1_: 0.0002275 s^-1^
~~~

~~~
k_2_: 0.195 s^-1^
~~~

~~~
k_3_: 0.2 s^-1^
~~~

~~~
k_4_: 0
~~~

~~~
k_5_: 0.00156 s^-1^
~~~

~~~
k_6_: 0.2 s^-1^
~~~

~~~
k_7_: 0.005 s^-1^
~~~

~~~
k_8_: 0.0001 s^-1^
~~~

~~~
k_9_: 0.005 s^-1^
~~~

~~~
k_10_: 0.0001 s^-1^
~~~

~~~
k_11_: 2.02e-06 molec^-1^ s^-1^
~~~

~~~
km_11_: 0.00252 s^-1^
~~~

~~~
k_12_: 0.05 s^-1^
~~~

~~~
k_13_: 4.5e-06 molec^-1^ s^-1^
~~~

~~~
km_13_: 5.36e-05 s^-1^
~~~

~~~
k_14_: 4e-05 s^-1^
~~~

~~~
k_k_: 5200 molec
~~~

~~~
k_s_: 1300 molec
~~~

~~~
n: 2
~~~

~~~
p: 3
~~~

~~~
Pconstitutive_ComK_: 1
~~~

~~~
Pconstitutive_ComS_: 1
~~~

~~~
P_ComK_: 1
~~~

~~~
P_ComS_: 1
~~~

~~~
M_T_: 520 molec
~~~

~~~
The simulation volume is Ω = 1 molec/nMolar.
~~~

The above equations produce molecular numbers. Concentrations are obtained by dividing the number of molecules by the simulation volume e.g. [ComK] = ComK /Ω. The units of concentration are therefore nanomolar. Dimensionless concentrations discussed in this work are obtained from the dimensioned concentrations using scaling factors:

[ComK]_dimensionless_ = [ComK] / Γ_ComK_ Γ_ComK_ = 2.6×10^4^

[ComS]_dimensionless_ = [ComS] / Γ_ComS_ Γ_ComS_ = 20.8

### Slowdown of translation in the canonical *Bacillus subtilis* circuit

The stochastic model is described at length in (2). The details of the model will not be reproduced here, for brevity. The slowdown of translation in the stochastic model as well as in its deterministic 6D infinite number of molecule limit model described in the preceding paragraph is implemented as follows:

~~~
k_3_ ← k_3_ × sdt_ComK_ Translation of mRNA_ComK
~~~

~~~
k_8_ ← k_8_ × sdt_ComK_ Degradation of ComK
~~~

~~~
k_11_ ← k_11_ × sdt_ComK_ ComK binding free MecA
~~~

~~~
km_11_ ← km_11_ × sdt_ComK_ MecA-ComK dissociation into MecA and ComK
~~~

~~~
k_12_ ← k_12_ × sdt_ComK_ Proteolytic degradation of MecA-bound ComK
~~~

~~~
k_6_ ← k_6_ × sdt_ComS_ Translation of mRNA_ComS
~~~

~~~
k_10_ ← k_10_ × sdt_ComS_ Degradation of ComS
~~~

~~~
k_13_ ← k_13_ × sdt_ComS_ ComS binding to free MecA
~~~

~~~
km_13_ ← km_13_ × sdt_ComS_ MecA-ComS dissociation into MecA and ComS
~~~

~~~
k_14_ ← k_14_ × sdt_ComS_ Proteolytic degradation of MecA-bound ComS
~~~

### Slowdown of translation in the inverted *Bacillus subtilis* circuit

The stochastic model is described at length in (2). The slowdown of translation in this model is implemented as follows:

~~~
k_5_ ← k_5_ × sdt Translation of mRNA ComK
~~~

~~~
k_6_ ← k_6_ × sdt Translation of mRNA MecA
~~~

~~~
k_10_ ← k_10_ × sdt Degradation of MecA
~~~

~~~
k_11_ ← k_11_ × sdt Degradation of ComK
~~~

### Scaling in the inverted *Bacillus subtilis* circuit

The scaling factor “sf” relating dimensioned to dimensionless concentrations ([]_dimensioned_ = []_dimensionless_ × sf) in the inverted *Bacillus subtilis* circuit is sf=25,000 (4).

### Stochastic simulations

Stochastic phase portraits in this work are occupancy diagrams (*i.e*. 2D histograms) generated by combining multiple (typically 200) stochastically independent histories binned over the range of the state variables. Each time history consists of an extended trajectory simulated by use of the Gillespie algorithm (10–16).

### Bifurcation Analysis

Bifurcation analyses were performed using Auto (17) and graphical interfaces XPP (18) and Oscill8 (19).

### Simulation Framework and Computing Platforms

All simulations were developed and performed using Matlab (The MathWorks, Natick, Massachusetts) on a Dell Intel i7-2640M dual-core Windows 7 laptop, and a Lenovo Intel i7-7700HQ quad-core laptop running Windows 10. Simulations required approximately ∼3000 hours (∼four full-time months) of integrated time over the course of a year.

## Discussion

In this work, we tackled the difficult problem of pinning down the respective roles of time-scale separation and of biological fluctuations in situations where these seemingly unrelated aspects of the dynamics conspire to affect the phenotype and might be individually or collectively harnessed by Evolution. We devised an effective method to remove fluctuations from the stochastic simulations without having to conduct a prohibitive number of discrete-event simulations to estimate the mean of their probability distributions. This approach yields a system of coupled ordinary differential equations from the set of discrete-event reactions. This born stochastic but now deterministic system conveniently continues to be endowed in the time scale separation underlying the stochastic dynamics. This approach simplifies studying the two conspiring aspects of the stochastic dynamics by separating their effects.

In the case of the canonical *Bacillus subtilis* regulation, we could then compare a 6D deterministic computation (with finite time-scale separation) to the 2D simplified view (with built-in infinite time-scale separation, the so-called *adiabatic approximation*). By performing a bifurcation analysis of the 6D system with respect to translation slowdown, we looked for and found effective (unequal) translation slowdown factors in the ComK and ComS dynamics that could then be applied to the stochastic dynamics. The anticipated result however, the presence of oscillations, did not have to immediately appear in a version of the stochastic dynamics now endowed with this slowed down translation. This is because stochasticity is, by design, not part of the 6D model and, yet, it could still be the source of the dynamical discrepancy. The anticipated oscillations did not, in fact, immediately occur in the slowed down translation version of the stochastic dynamics. But by reducing the amount of biochemical noise in the system, we then discovered that this is because of the conspiring effect of stochasticity. Hence, only by further reducing the intrinsic noise in the system, did the expected oscillations finally occur (Figure 5, panel c)). Note that reducing stochasticity alone in the system (leaving time-scale separation intact) was determined to be insufficient to resolve the dynamical discrepancy (data omitted for brevity).

In the case of the inverted *Bacillus subtilis* regulation, we showed that oscillations that should have already vanished from an infinite time-scale separation perspective persisted only due to remaining finite time-scale separation in the stochastic system. By slowing down translation on both ComK and MecA (here, by the same factor) in the discrete event simulation, oscillations expected to have vanished did, in fact, then vanish (compare Figure 8 panel b) and d). Additionally, we showed that stochasticity alone could not resolve the dynamical mismatch.

### Is stochasticity truly an orthogonal evolutionary handle to time-scale separation?

It is conceivable that Evolution can adjust relevant constituent molecular numbers to affect stochasticity. It is also similarly conceivable that Evolution can independently affect time scale separation by adjusting relevant rate parameters (*i.e*. fine tuning binding constants (20–24)). However, are these two evolutionary handles necessarily orthogonal? Evolution has no effect on basic physics; we note that cell size couples these two aspects. Physically larger cells have more molecules in a concomitant proportionately bigger volume so they are expected to be less noisy. This expectation has been confirmed experimentally by the fusion of two bacterial cells together (6). However, because transcription and translation are physically separate non-co-located processes, physically larger cells would also be expected to have, on average, more time scale separation. Therefore, for larger cells, physics hints that noise would be reduced and time-scale separation would be increased. These are exactly the conditions we have observed in this work that led to phenotype breaking in the canonical circuit: increased time-scale separation and reduced noise caused a dynamical regime change from biologically desirable excitability to undesirable oscillations.

## Conclusions

Time-scale separation is controlled by relationships between kinetic parameters in the mRNA and protein system whereas stochasticity is controlled by the number of molecules in the system. Therefore, these two dynamical aspects offer accessible “handles” that Evolution may be able to adjust independently to affect phenotype: e.g. as studied here, cause or avoid oscillations. In this work, for one evolutionary realized canonical bacterium and one hypothetical re-engineered mutant, we specifically point out how these handles can be used to affect the range over which oscillations occur. Elsewhere (2) it has already been shown that oscillations are detrimental to the competence phenotype. Here we show that biologically undesirable oscillatory regimes are, in fact, present in both versions of the control circuits (canonical and inverted). Moreover, for the canonical competence circuit, onset of possibly phenotypically catastrophic large oscillations can be guarded against using both time-scale separation and fluctuation levels originating in the paucity of some of the molecular components in the system. On the other hand, for the inverted competence circuit, time-scale separation alone is sufficient to guard against unwanted oscillations.

## Contributions

LH and ZY Performed 6D bifurcation analysis.

MT Devised the research. Performed bifurcation analysis. Performed stochastic and deterministic simulations. Wrote the paper.

## Acknowledgements

MT would like to thank Chen Ling and Wiegang Sun for their hospitality during a stay at Hangzhou Dianzi University in Hangzhou and Jinzhi Lei for his hospitality during a stay at Tsinghua University in Beijing where some aspects of this research were developed.

## Reference List

1. Eldar, A. and Elowitz, M. B. (2010) Nature 467, 167–173

2. Cagatay, T., Turcotte, M., Elowitz, M. B., Garcia-Ojalvo, J., and Suel, G. M. (2009) Cell 139, 512–522

3. Schultz, D., Ben Jacob, E., Onuchic, J. N., and Wolynes, P. G. (2007) Proceedings of the National Academy of Sciences 104, 17582–17587

4. Turcotte, M., Garcia-Ojalvo, J., and Suel, G. M. (2008) Proceedings of the National Academy of Sciences 105, 15732–15737

5. Süel, G. M., Garcia-Ojalvo, J., Liberman, L. M., and Elowitz, M. B. (2006) Nature 440, 545–550

6. Suel, G. M., Kulkarni, R. P., Dworkin, J., Garcia-Ojalvo, J., and Elowitz, M. B. (2007) Science 315, 1716–1719

7. Dubnau, D. (1991) Genetic competence in Bacillus subtilis.

8. Dubnau, D. (1991) Molecular Microbiology 5, 11–18

9. Dubnau, D. (1999) Annu. Rev. Microbiol. 53, 217–244

10. Gillespie, D. T. (1976) Journal of Computational Physics 22, 403–434

11. Gillespie, D. T. (1977) J Phys Chem 81, 2340–2361

12. Gillespie, D. T. (2001) J Chem Phys 115, 1716–1733

13. Gillespie, D. T. (1976) Journal of Computational Physics 22, 403–434

14. Gillespie, D. T. (1977) The Journal of Physical Chemistry 81, 2340–2361

15. Gillespie, D. T. (1991) Markov Processes: An Introduction for Physical Scientists, Academic Press,

16. Gillespie, D. T. (2007) Annu. Rev. Phys. Chem. 35–55

17. Doedel, E. J. (2008) AUTO: Software for Continuation and Bifurcation Problems in Ordinary Differential Equations.

18. Ermentrout, B. (2011) XPP-Aut (http://www.math.pitt.edu/~bard/xpp/xpp.html).

19. Conrad, E. D. (2011) Oscill8 (http://sourceforge.net/projects/oscill8/).

20. Levy, E. D. (2010) Journal of Molecular Biology 403, 660–670

21. Kastritis, P. L. and Bonvin, A. M. (2013) Current Opinion in Structural Biology 23, 868–877

22. Kastritis, P. L., Rodrigues, J. P. G. L., Folkers, G. E., Boelens, R., and Bonvin, A. M. J. J. (2014) Journal of Molecular Biology 426, 2632–2652

23. Rosanova, A., Colliva, A., Osella, M., and Caselle, M. (2017) Scientific Reports 7, 7596

24. Echave, J. and Wilke, C. O. (2017) Annu. Rev. Biophys. 46, 85–103

